# Anti-Sense Oligonucleotide as a Therapeutic for Synucleinopathies: Pharmacokinetic, Safety and Efficacy Evaluation

**DOI:** 10.1101/2025.05.01.651722

**Authors:** Rijwan U. Ahammad, Brian Spencer, Bao Quach, Sahar Salehi, Robert A. Rissman

**Author notes:** **Corresponding Author:** Robert A. Rissman, PhD, Alzheimer’s Therapeutic Research Institute, Keck School of Medicine of University of Southern California, 9880 Mesa Rim Road, San Diego, CA, 92121, USA.

## Abstract

Effective blood-brain barrier (BBB) penetration is a significant challenge for antisense oligonucleotide (ASO) therapies targeting neurodegenerative diseases. We utilized a peptide (ApoB^11^) mediated transport delivery of an ASO to the CNS following systemic delivery to reduce expression of targeted transcripts for neurodegenerative diseases. This study evaluates the pharmacokinetics, CNS penetration, and therapeutic efficacy of ApoB^11^:2’-OMe ASO-α-Syn, an ASO for α-synuclein (α-Syn) suppression in synucleinopathies. After a single intraperitoneal (IP) injection (2 mg/kg) in C57BL/6 mice, ApoB^11^:ASO-α-Syn showed robust brain penetration, reaching peak concentrations (C_max_ = 0.14 nMol/mg) at 1.5 hours and an extended brain half-life (t_1/2_ = 646.2 hours), indicating prolonged CNS retention. Immunofluorescence confirmed widespread uptake in neurons and endothelial cells. The ASO also accumulated in the liver (C_max_ = 419.5 nMol/mg, t_1/2_ = 104.9 hours), consistent with receptor-mediated uptake. Acute and subacute toxicity studies revealed no systemic toxicity at the highest non-lethal dose (32 mg/kg). In a mouse model of dementia with lewy body (DLB) mice overexpressing human α-Syn, ApoB^11^:ASO-α-Syn reduced α-Syn mRNA and protein levels in the hippocampus and cortex by ∼50% at 16 mg/kg. These results demonstrate that ApoB^11^ is an effective ASO carrier for CNS delivery, supporting its potential as a therapeutic strategy for synucleinopathies.

**Graphical Abstract:** 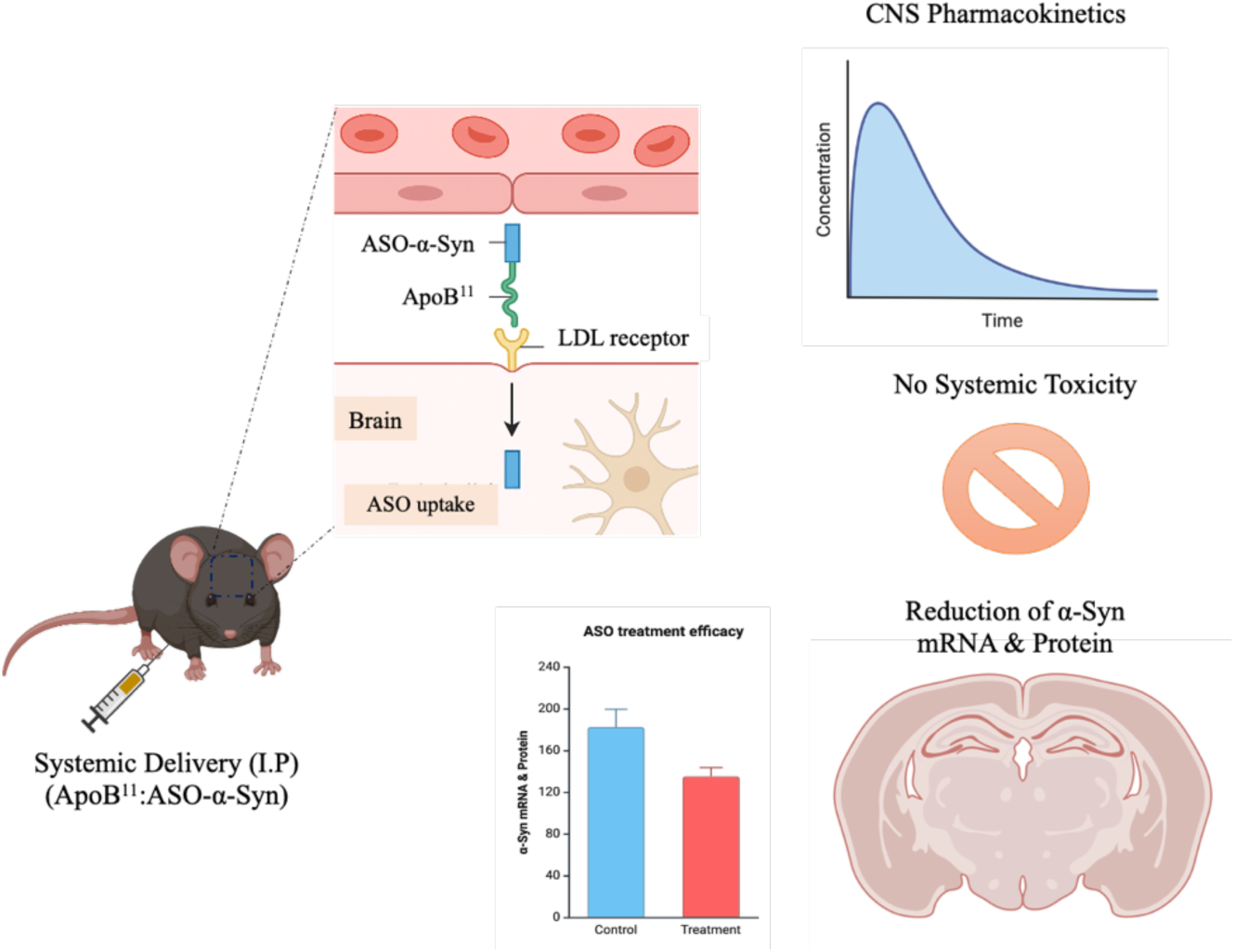

## INTRODUCTION

Antisense oligonucleotides (ASOs) represent an innovative therapeutic platform for treating central nervous system (CNS) diseases by enabling the selective suppression of disease driving genes. These synthetic, single-stranded oligonucleotides bind to complementary RNA sequences, leading to degradation of the target transcript or inhibition of its translation.^1,2^ ASOs offer several advantages over other gene-targeting approaches, including reversibility, dose-dependence, and a low risk of insertional mutagenesis.^3^ Additionally, chemical modifications can enhance their stability, reduce immune activation, and improve *in vivo* efficacy.^4^ Given these advantages, ASOs have emerged as a versatile and promising tool for modulating gene expression in the CNS.

Compared to RNA interference (RNAi), which requires cytoplasmic delivery and depends on the endogenous RISC complex, ASOs can act in both the nucleus and cytoplasm, and they do not saturate endogenous RNA processing pathways.^5^ This enables more predictable and dose-controllable gene knockdown. Similarly, unlike adeno-associated virus (AAV)-mediated gene therapies, which induce long-term transgene expression and raise concerns about immune response and genomic integration, ASOs do not integrate into the genome and offer transient, titratable effects, allowing greater flexibility in treatment regimens.^6,7^ These properties make ASOs especially well-suited for targeting genes involved in progressive and region-specific CNS disorders.

Despite their promise, a major limitation of ASO therapies for CNS disorders is effective delivery across the blood-brain barrier (BBB). Current clinical applications typically rely on intrathecal or intracerebroventricular administration routes that are invasive and not ideal for long-term treatment.^8,9^ Systemic delivery is a more practical and scalable approach, but faces challenges including poor BBB permeability, off-target exposure, and inefficient uptake into neurons.^10,11^ Thus, strategies to enhance brain delivery of ASOs are crucial for advancing their clinical translation.

To overcome these delivery barriers, ligand-mediated strategies have gained increasing interest. Among them, transport peptides such as ApoB^11^, a sequence derived from apolipoprotein B show promise for facilitating receptor-mediated transcytosis across the BBB. ApoB^11^ binds to low-density lipoprotein (LDL) receptors, promoting uptake into the brain parenchyma.^12-14^ When conjugated to ASOs, these peptides can enable non-invasive, systemic delivery with improved biodistribution, offering a potential paradigm shift in CNS drug delivery.

In this study, we used an ApoB^11^ conjugated ASO targeting human α-synuclein (α-Syn) mRNA to evaluate its pharmacokinetics (PK), biodistribution, and pharmacodynamic (PD) efficacy following systemic administration. α-Syn is a presynaptic neuronal protein that plays a central role in synucleinopathies such as Parkinson’s disease (PD), dementia with Lewy bodies (DLB), and multiple system atrophy (MSA).^15^ In these diseases, α-Syn undergoes pathological misfolding and aggregation, forming oligomers and fibrils that disrupt cellular homeostasis and promote neurodegeneration.^16^

Importantly, the accumulation of α-Syn is not only a hallmark of these disorders but also correlates with disease progression and symptom severity, making it a compelling therapeutic target. Reducing α-Syn expression has been shown to attenuate pathology and improve outcomes in various preclinical models,^17,18^ further supporting its relevance for evaluating ASO efficacy. Thus, α-Syn serves as both a mechanistically relevant target and a quantitative readout for assessing CNS delivery and gene silencing effects.

Here, we assessed the *in vivo* distribution, target engagement, and therapeutic impact of systemically administered ApoB^11^:ASO-α-Syn conjugates in a transgenic DLB mouse model overexpressing human α-Syn. We specifically examined whether this approach enables effective BBB crossing and gene knockdown in disease relevant regions such as the cortex and hippocampus. This work aims to provide a comprehensive PK/PD profile of a systemically delivered, CNS-targeting ASO and contribute to the development of minimally invasive, disease-modifying therapies for neurodegenerative disorders.

## RESULTS

### Pharmacokinetics and CNS Penetration of ApoB^11^:ASO-α-Syn

Achieving efficient BBB penetration is a key hurdle for systemically administered ASO therapeutics. In this study, we evaluated the pharmacokinetic (PK) profile and tissue biodistribution of ApoB^11^:ASO-α-Syn, a BBB-permeable ASO, following a single intraperitoneal (IP) administration (2 mg/kg) in male C57BL/6 mice. ASO concentrations in plasma and tissues were quantified using a validated ASO bioassay, and concentration-time profiles were analyzed to assess systemic absorption, clearance, and organ-specific uptake.

ApoB^11^:ASO-α-Syn demonstrated robust brain penetration, reaching peak ASO concentrations (C_max_) of 0.14 nMol/mg at 1.5 hours post-injection (T_max_) (Figure 1A, 1F). Notably, the extended brain half-life (t_1/2_ = 646.2 hours) indicates slow clearance and sustained retention, suggesting prolonged engagement with CNS targets. This retention profile is consistent with ApoB^11^-mediated transcytosis across the BBB,^13^ supporting its role as an effective ASO delivery platform for neurological diseases.

**Figure 1:**
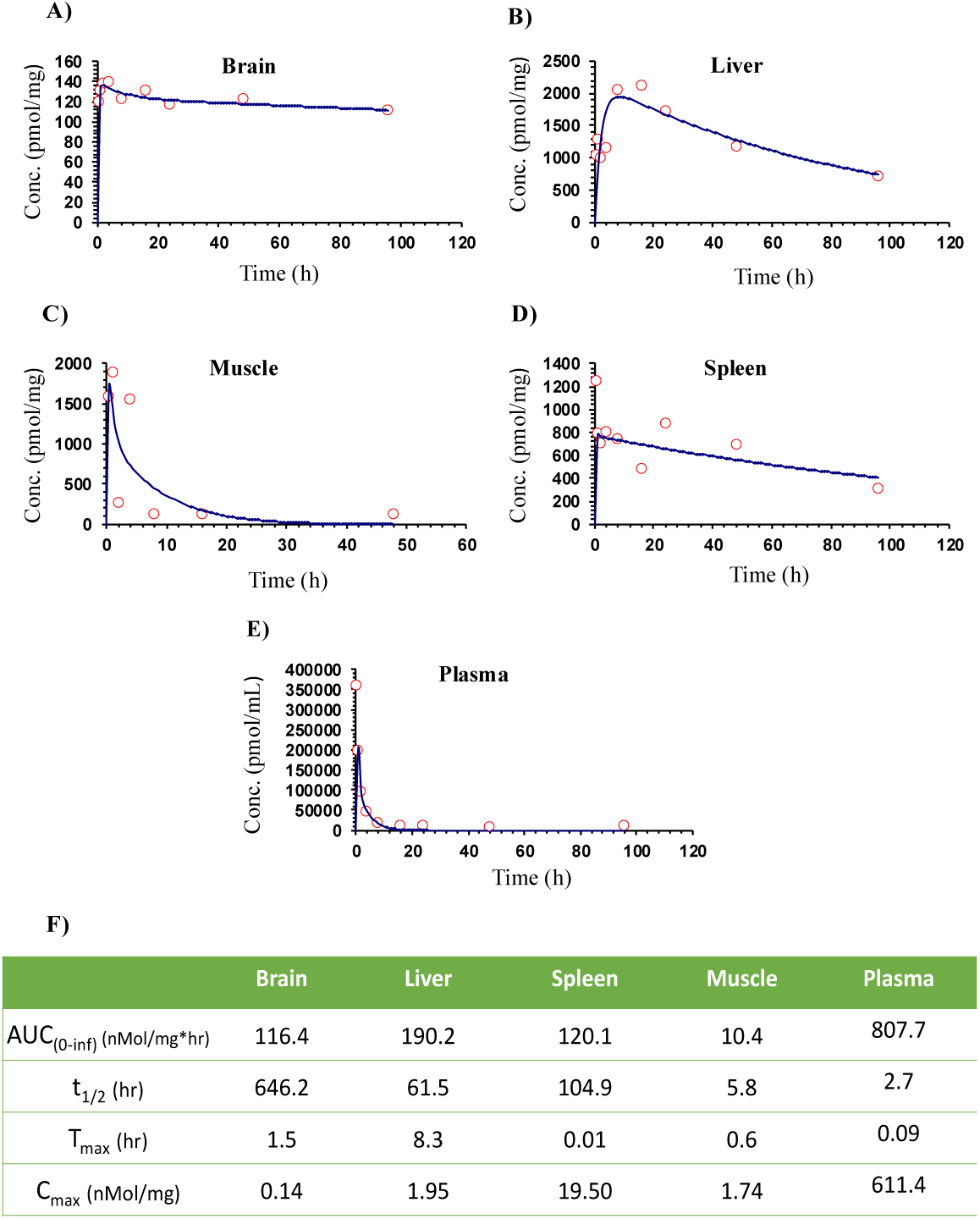
Pharmacokinetic profiling and brain penetration of ApoB^11^:ASO-α-Syn following systemic administration. (A–E) Concentration–time profiles of ApoB^11^:ASO-α-Syn in plasma, liver, spleen, skeletal muscle, and brain of male C57BL/6 mice (n = 3 per timepoint) following a single intraperitoneal injection (2 mg/kg). ASO concentrations were quantified at multiple timepoints (up to 120 hours) using a hybridization-based ASO bioassay. (F) Summary table showing pharmacokinetic parameters of ApoB^11^:ASO-α-Syn following systemic administration modeled with the Microsoft Excel plugin pkpdtools.

Beyond CNS uptake, ApoB^11^:ASO-α-Syn accumulates in the liver significantly influencing the systemic pharmacokinetics and tissue biodistribution of the ASO (Figure 1B,1F). Plasma ASO levels peaked rapidly (C_max_ = 611.4 nMol/mg, T_max_ = 0.09 hours) following IP administration, with a moderate systemic half-life (t_1/2_ = 2.7 hours) and an AUC_0-οο_ of 807.7 nMol·h/mg, indicative of efficient absorption and clearance (Figure 1E, 1F). Liver exhibited the highest ASO accumulation (C_max_ = 419.5 nMol/mg, T_max_ = 8.3 hours), with sustained retention (t_1/2_ = 104.9 hours), suggesting receptor-mediated uptake^19^. This prolonged hepatic half-life aligns with known ApoB^11^-mediated ASO internalization and suggests potential hepatic involvement in ASO clearance. The spleen showed moderate uptake (C_max_ = 120.1 nMol/mg, T_max_ = 0.6 hours, t_1/2_ = 5.8 hours), supporting its role in systemic ASO distribution and clearance (Figure 1D, 1F). In contrast, ASO uptake in muscle was minimal (C_max_ = 1.74 nMol/mg, T_max_ = 0.6 hours), with a rapid clearance profile (t_1/2_ = 5.8 hours, AUC_₀–∞_ = 10.4 nMol·h/mg), indicating limited peripheral muscle retention (Figure 1C, 1F).

### Biodistribution of ApoB^11^:ASO-α-Syn in Brain Tissue Following Systemic Administration

Following pharmacokinetic (PK) analysis demonstrating the presence of ApoB^11^:ASO-α-Syn in the CNS, we next examined its cellular distribution within different brain regions and CNS cell types. Given the sustained ASO-α-Syn retention in the brain, we hypothesized that it localizes to neuronal and non-neuronal compartments, potentially influencing α-synuclein pathology across multiple cell populations. To assess this, wild-type (WT; C57BL/6) mice received a single intraperitoneal (IP) injection of biotinylated ASO-α-Syn (2 mg/kg) and were sacrificed 24 hours post-injection. Brain tissue sections were analyzed via immunofluorescence staining, co-labeling for neuronal (NeuN), astrocytic (GFAP), microglial (Iba1), oligodendrocytic (Olig2), and endothelial (Lectin) markers to determine ASO localization (Figure 2A, 2B).

**Figure 2:**
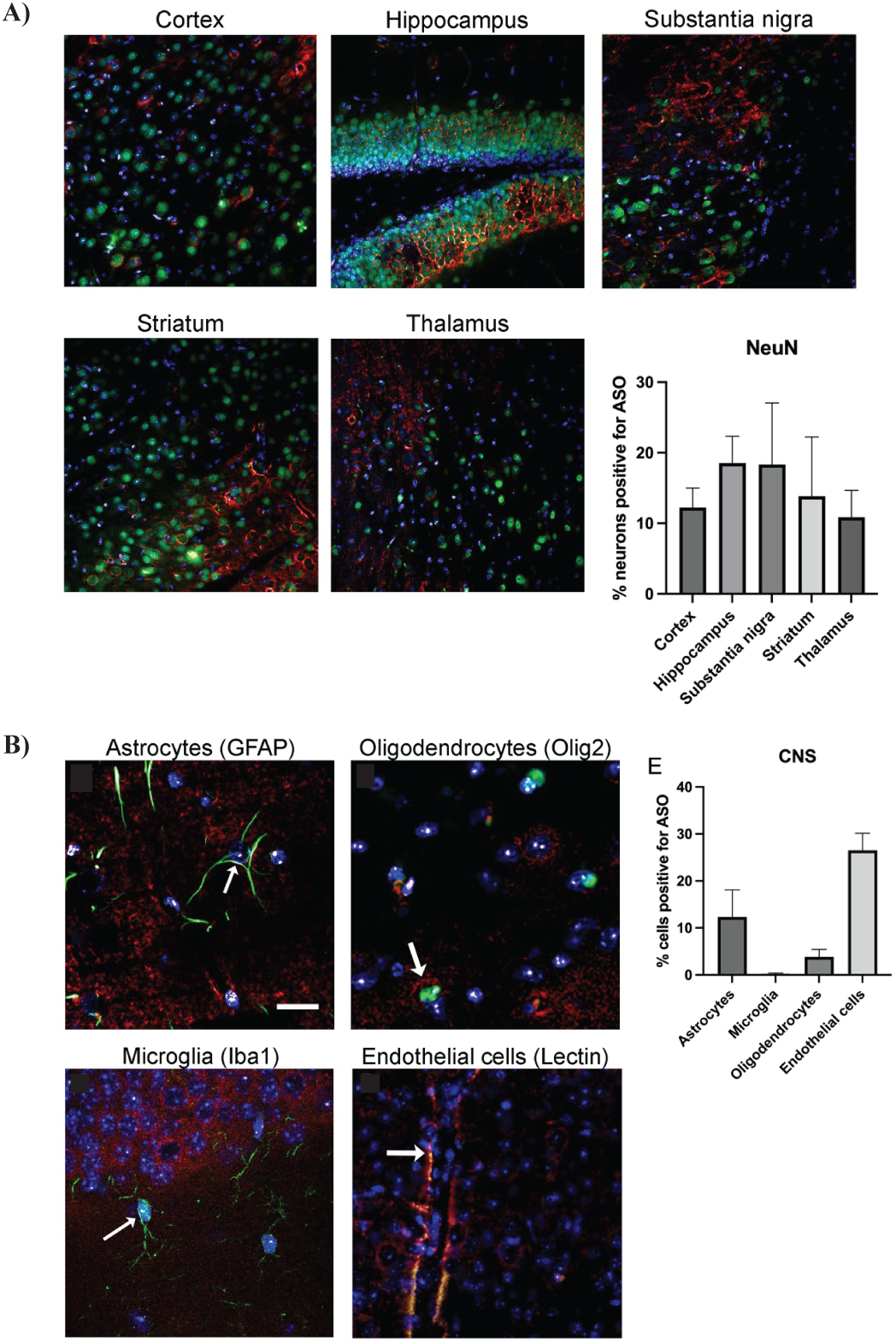
Cellular distribution of ApoB^11^:ASO-α-Syn in the central nervous system. Wild-type C57BL/6 mice (n = 3) received intraperitoneal injection of biotin-labeled α-Syn ASO conjugated to ApoB^11^ (2 mg/kg) to assess CNS delivery. Biotin-tagged ASO was used to visualize drug localization in brain tissue. (A) Representative immunofluorescence images showing ASO-α-Syn localization (green) in the cortex, hippocampus, substantia nigra, striatum, and thalamus. Neuronal marker NeuN (red) and nuclear stain DAPI (blue) are shown. Quantification of NeuN⁺ neurons positive for ASO-α-Syn reveals highest uptake in substantia nigra (18.5%) and hippocampus (18%), followed by striatum (14.5%), cortex (12%), and thalamus (9%). (B) Immunofluorescence staining for ASO-α-Syn (green) in CNS glial and endothelial cells, including astrocytes (GFAP, red), oligodendrocytes (Olig2, red), microglia (Iba1, red), and endothelial cells (Lectin, red). White arrows indicate ASO-positive cells. Quantification of ASO-positive cells per cell type shows highest uptake in endothelial cells (26%), followed by astrocytes (12%) and oligodendrocytes (4.5%). Microglial uptake was minimal (0.5%). Data are shown as mean ± SD

Fluorescence imaging revealed widespread ASO-α-Syn uptake across multiple brain regions, with the highest neuronal accumulation observed in the substantia nigra (18.5%), hippocampus (18%), striatum (14.5%), and cortex (12%), while the thalamus exhibited the lowest uptake (9%) (Figure 2A). Within neuronal populations, ASO-α-Syn was detected within neuronal cell bodies and their surrounding neuropil, suggesting active cellular uptake rather than passive diffusion. Among non-neuronal cell types, endothelial cells (26%) showed the highest ASO-α-Syn signal. Astrocytes exhibited moderate ASO uptake (12%), whereas oligodendrocytes showed lower levels (4.5%). Microglial uptake was minimal (0.5%), within the analyzed timeframe (Figure 2B).

Beyond the CNS, ASO biodistribution was assessed in peripheral organs, including the liver, spleen, and muscle. Immunofluorescence staining revealed significant ASO accumulation in hepatocytes, consistent with the liver’s high uptake capacity (Figure S1A). The spleen exhibited moderate ASO signal, particularly in immune cell-rich regions, supporting its role in systemic ASO distribution and clearance (Figure S1B). In contrast, skeletal muscle showed minimal ASO uptake, suggesting limited penetration into muscle fibers (Figure S1C). These findings indicate that ApoB^11^-ASO -α-Syn, efficiently distributes across the CNS and periphery, with predominant localization in neurons, endothelial cells, and hepatocytes.

ApoB^11^:ASO-α-Syn is Well-Tolerated with No Observed Acute or Subacute Toxicity

To determine the maximum tolerated dose (MTD) of ApoB^11^:ASO-α-Syn, an acute toxicity study was conducted in wild-type (WT) C57BL/6 mice using intraperitoneal (IP) administration of increasing doses (32, 50, and 75 mg/kg). Each experimental group consisted of equal numbers of male and female mice (n=8 per group; 4 males and 4 females). A dose of 75 mg/kg was lethal in all animals following the first injection, while 50 mg/kg resulted in progressive mortality, with all mice succumbing by the third injection. In contrast, mice administered 32 mg/kg showed no mortality or signs of acute toxicity, establishing this as the highest non-lethal dose (Table S1).

To evaluate potential long-term toxic effects, we monitored the overall health and growth response of mice receiving weekly injections of 32 mg/kg ApoB^11^:ASO-α-Syn for four weeks, followed by a four-week recovery period. Male mice initially experienced a slight decrease in body weight during the treatment phase; however, their weight gradually recovered over time, aligning with the control group by the end of the study. In contrast, female mice showed no significant weight differences between treatment and control groups throughout the study. No behavioral abnormalities or adverse health effects were observed in any group (Figure S2). These findings suggest ApoB^11^:ASO-α-Syn repeated administration at the maximum tolerated dose does not induce long-term systemic toxicity or negatively impact normal physiological functions.

To further evaluate the safety profile ApoB^11^:ASO-α-Syn, histopathological analyses were conducted on tissues from mice administered either 32 mg/kg or 10 mg/kg ApoB^11^:ASO-α-Syn, alongside saline controls, over a four-week period (Figure 3A). Hematoxylin and eosin (H&E) staining of major peripheral organs, including the liver, kidney, spleen, heart, lung, skeletal muscle, and spinal cord, revealed no evidence of necrosis, inflammation, or cellular abnormalities (Table S2). Additionally, representative H&E-stained images of brain tissues, including the hippocampus and cortex, from high-dose (32 mg/kg) and control groups confirmed the absence of any histopathological abnormalities (Figure 3B). In the analysis of major organs, no histological changes were observed in any of the tissues from the treated mice when compared to saline controls (Figure 3C). These results suggest that ApoB^11^:ASO-α-Syn, at doses of 32 mg/kg and 10 mg/kg (Data not shown), does not induce significant toxicity or histopathological alterations in the peripheral organs or brain tissues of treated mice.

**Figure 3:**
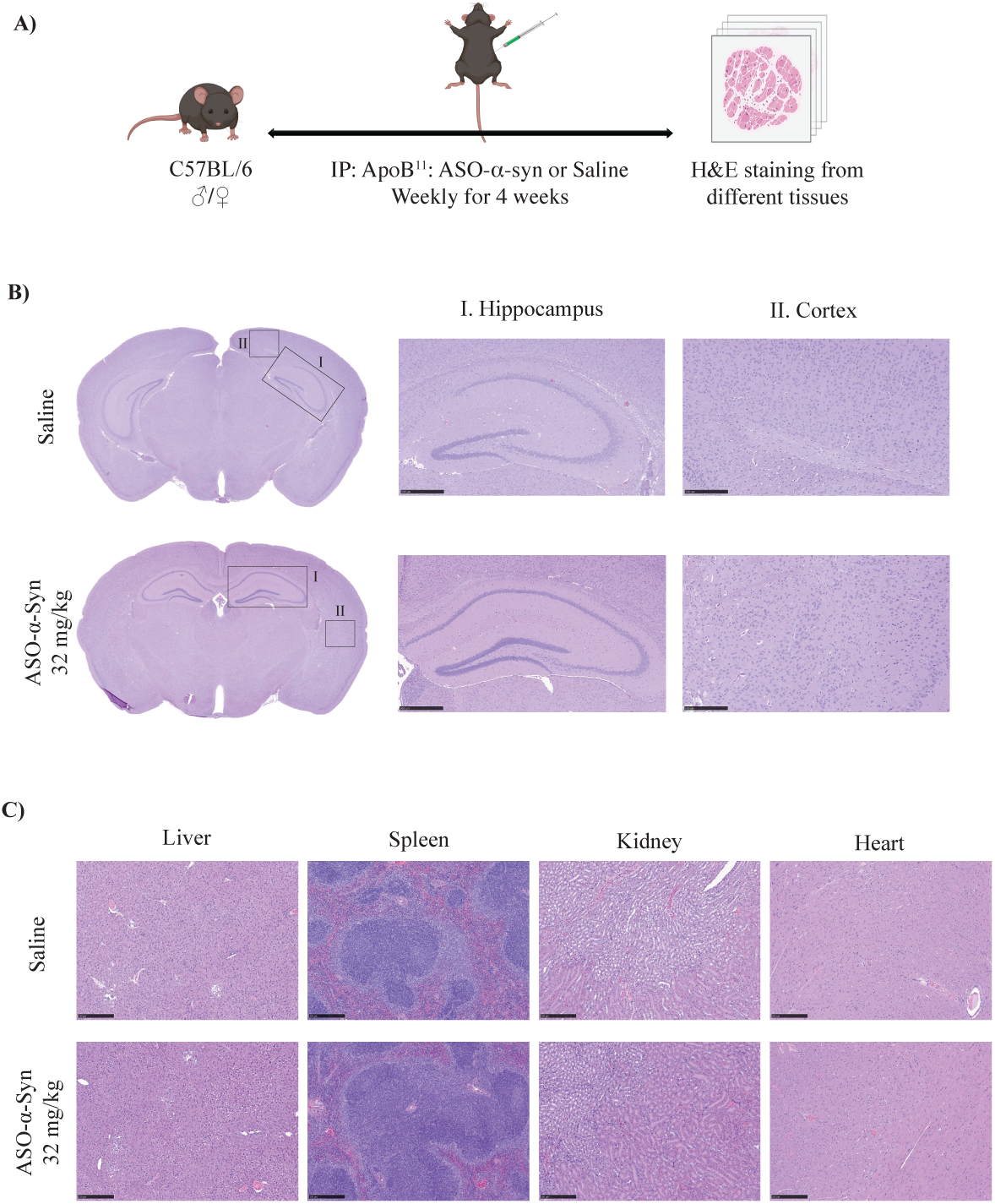
Histopathological assessment following repeated ApoB^11^:ASO-α-Syn administration. (A) Study design outlining repeated high-dose administration of ApoB^11^:ASO-α-Syn. (B) Representative H&E-stained sections of brain (hippocampus and cortex) from saline-treated and high-dose (32 mg/kg) treated mice show no evidence of necrosis, inflammation, or cellular degeneration. (C) Representative H&E-stained images of peripheral organs (liver, spleen, kidney, heart) reveal no pathological abnormalities. Scale bars: hippocampus (5×, 500 µm), cortex (10×, 250 µm), other organs (10×, 250 µm).

Dose-Dependent Reduction of α-Syn by ApoB^11^:ASO-α-Syn in D-Line Transgenic Mice

D-line transgenic mice, which overexpress human α-synuclein (hu-αSyn), were selected for this study due to their well-characterized synucleinopathy, making them a suitable model for evaluating antisense oligonucleotide (ASO) mediated knockdown strategies. This model exhibits progressive accumulation of hu-αSyn in key brain regions, including the hippocampus and cortex, mirroring pathophysiological features observed in synucleinopathies such as Parkinson’s disease and dementia with Lewy bodies.^20,21^ To determine the efficacy of ApoB^11^:ASO-α-Syn in reducing hu-αSyn expression, we administered a single bolus (0.5, 2, 8, 16, and 32 mg/kg) injection (IP) and evaluated changes in hu-αSyn mRNA and protein levels 7 days after injection (Figure 4A).

**Figure 4:**
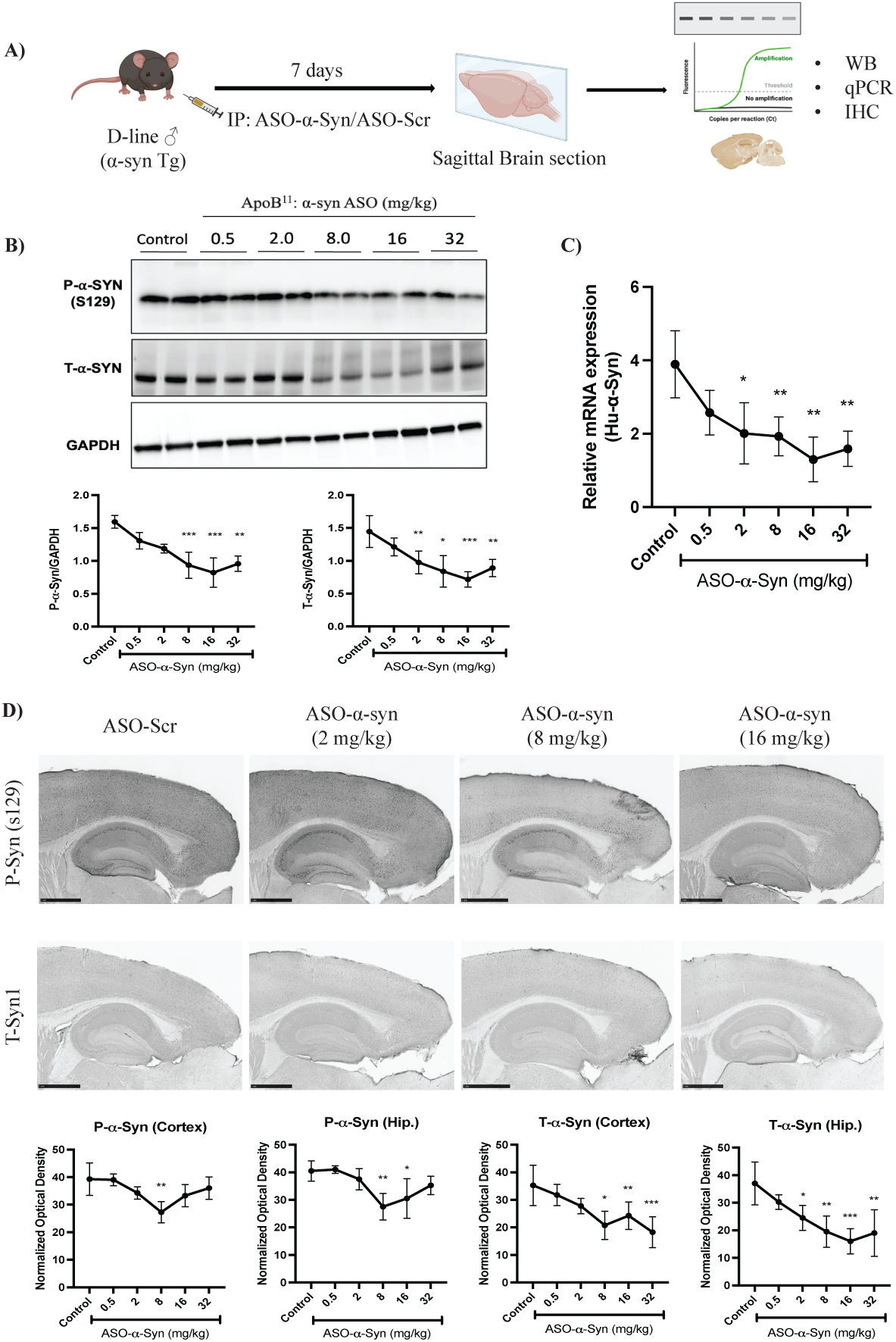
Dose-dependent reduction of α-Syn pathology by ApoB^11^:ASO-α-Syn in D-line mice. (A) Shows experimental design. Human α-Syn transgenic (D-line) mice received a single intraperitoneal dose of ApoB^11^:ASO-α-Syn (0.5, 2.0, 8.0, 16.0, or 32.0 mg/kg). Brains were collected 7 days later for molecular and histological analyses. (B) Representative western blot analysis of phosphorylated α-synuclein (P-α-Syn, Ser129), total α-synuclein (T-α-Syn), and GAPDH. Quantification of P-α-Syn (left) and total T-α-Syn (right), normalized to GAPDH, is shown in the bottom panel. (C) Relative expression of human α-synuclein (huSyn) mRNA in brain tissue measured by qRT-PCR, normalized to β-actin. (D) Immunohistochemistry images showing P-α-Syn and T-α-Syn in hippocampus and cortex. Quantification of immunoreactive area for P-α-Syn (left) and T-α-Syn (right). Scale bar = 500 µm. Data are mean ± SD (n = 4 per group). One-way ANOVA followed by Dunnett’s post hoc test; *p < 0.05, **p < 0.01 vs. control.

Western blot (WB) analysis of brain lysates, primarily from the hippocampus and cortex, revealed a dose-dependent reduction in total α-Syn (T-α-Syn) and phosphorylated α-Syn at serine 129 (P-α-Syn S129) levels (Figure 4B). While lower doses (0.5 and 2 mg/kg) had minimal effect, significant suppression of ∼50% was observed at doses of 8 mg/kg and above (p < 0.01). The maximal reduction occurred at 16 mg/kg, with no further decrease at 32 mg/kg, suggesting a plateau effect. These findings indicate that ApoB^11^:ASO-α-Syn effectively reduces α-Syn protein levels in a dose-dependent manner in the hippocampus and cortex.

Quantitative RT-PCR (qRT-PCR) analysis of hu-α-Syn mRNA, corroborated the WB findings, showing a corresponding reduction in transcript levels in brain (Figure 4C). The decrease in hu-αSyn mRNA was significant at 8 mg/kg and peaked at 16 mg/kg, consistent with the protein-level data. This alignment between mRNA and protein reduction confirms that ApoB^11^:ASO-α-Syn effectively suppresses hu-αSyn expression at the transcriptional level.

Immunohistochemistry (IHC) further validated these results by demonstrating reduced accumulation of both P-α-Syn and T-α-Syn in the hippocampus and cortex (Figure 4D). Representative images were shown for selected doses (2, 8, and 16 mg/kg) to illustrate the dose-dependent reduction in P-α-Syn and T-α-Syn staining, while quantitative analysis was provided for all treatment groups, including 0.5 and 32 mg/kg. The reduction in both P-α-Syn and T-α-Syn staining intensity at 8 mg/kg and above was consistent with WB and qRT-PCR results, reinforcing the robust efficacy of ApoB^11^:ASO-α-Syn in lowering pathological α-Syn levels in key brain regions.

### Time-Dependent Reduction of α-Syn by ApoB^11^:ASO-α-Syn in D-Line Transgenic Mice

To evaluate the sustained efficacy and pharmacokinetics of ApoB^11^:ASO-α-Syn, a single intraperitoneal (IP) injection (2 mg/kg) was administered to D-line transgenic mice, and α-Syn levels were assessed at different time points post-injection (Day 1, 7, 14, 21, 28, and 35) (Figure 5). Western blot analysis of brain lysates demonstrated a progressive decline in both total α-Syn (T-α-Syn) and phosphorylated αSyn at serine 129 (P-α-Syn S129) (Figure 5A). The reduction was significant by Day 7 (∼30%), reached a maximum at Day 14 (∼60%), and persisted through Day 21 before gradually returning toward baseline by Day 28 & 35, likely due to ASO clearance. These findings suggest that a single-dose administration of ApoB^11^:ASO-α-Syn leads to prolonged α-Syn suppression, with peak efficacy occurring between Day 7 and Day 21.

**Figure 5:**
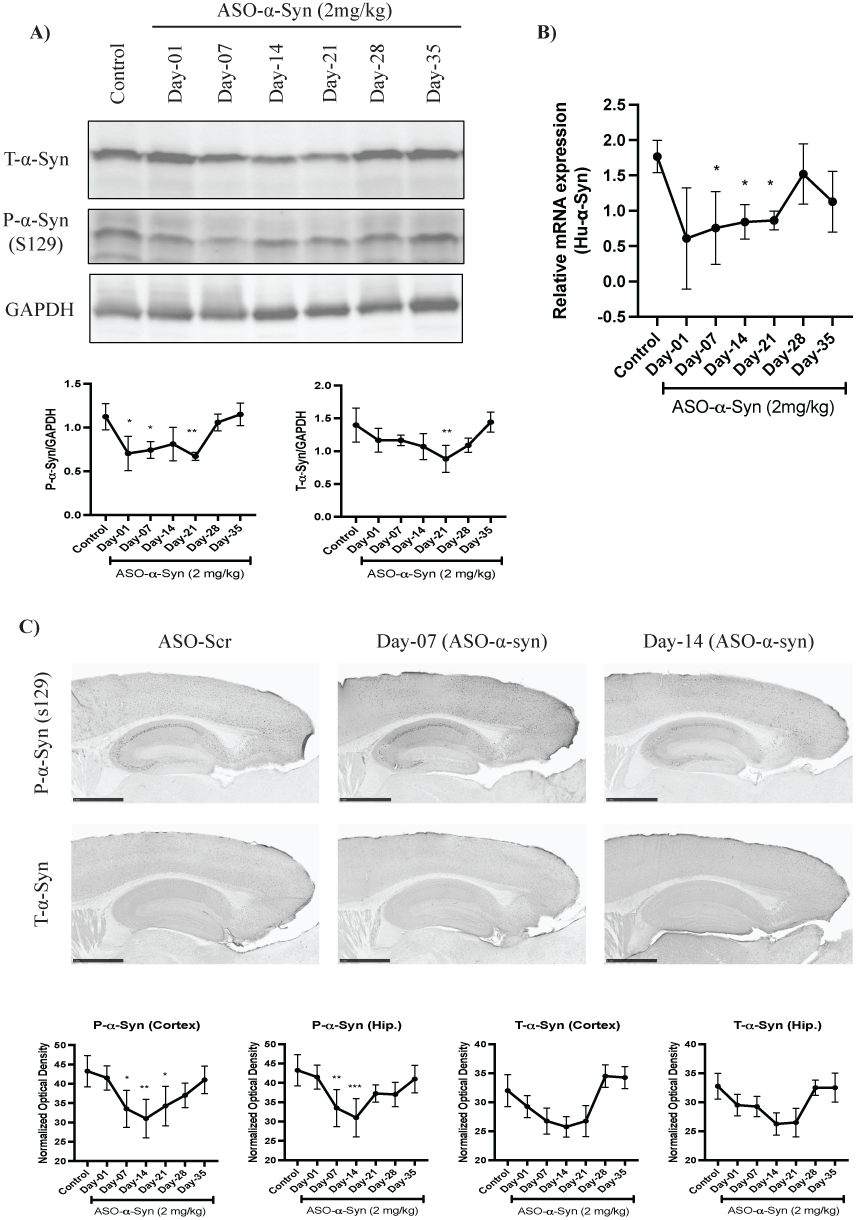
Time-dependent suppression of α-Syn pathology following ApoB^11^:ASO-α-Syn treatment in D-line mice. D-line transgenic mice received a single intraperitoneal dose of ApoB^11^:ASO-α-Syn (2 mg/kg) and were sacrificed at various timepoints for molecular and histological analyses. (A) Representative western blot images of phosphorylated α-synuclein (P-α-Syn, Ser129), total α-synuclein (T-α-Syn), and GAPDH from brain lysates of mice treated with vehicle, control ASO, or ApoB^11^:ASO-α-Syn. Quantification of P-α-Syn (top) and T-α-Syn (bottom), normalized to GAPDH, is shown below. (B) Relative expression of human α-synuclein (huSyn) mRNA in brain tissue assessed by qRT-PCR, normalized to β-actin. (C) Representative immunohistochemistry images showing P-α-Syn and T-α-Syn in hippocampus and cortex across treatment groups. Quantification of immunoreactive area for P-α-Syn (left) and T-α-Syn (right) is shown. Scale bar = 500 µm. Data are presented as mean ± SD (n = 4 per group). Statistical significance was determined by one-way ANOVA followed by Tukey’s post hoc test (*p < 0.05, **p < 0.01, ***p < 0.001).

The qRT-PCR analysis corroborated the protein-level data, revealing a similar trend in hu-αSyn mRNA expression. A maximal suppression of ∼55% was observed Day 14, with a gradual increase by Day 28, confirming that the ASO acts at the transcriptional level to reduce α-Syn expression (Figure 5B). Immunohistochemistry (IHC) further supported these findings, showing a gradual decrease in αSyn immunoreactivity in hippocampal and cortical sections, with the most significant reductions at Day 07 and Day 14. By Day 28, a partial recovery in αSyn staining was observed, reinforcing the transient nature of ASO-mediated knockdown (Figure 5C). Together, these results highlight ApoB^11^:ASO-α-Syn’s sustained efficacy and pharmacokinetics, positioning it as a promising long-acting therapeutic strategy for synucleinopathies.

## DISCUSSION

This study provides compelling evidence that ApoB^11^:ASO-α-Syn holds significant potential as a therapeutic strategy for synucleinopathies, particularly in targeting α-Syn expression. The efficient BBB penetration, robust neuronal uptake, and prolonged retention in the central nervous system (CNS) are essential factors for achieving sustained therapeutic engagement of neuronal targets, particularly in chronic neurodegenerative diseases such as Parkinson’s disease (PD) and dementia with Lewy bodies (DLB). These findings align with prior reports of ligand-mediated antisense oligonucleotide (ASO) transport across the BBB,^4,22,23^ and underscore the unique advantages of ApoB^11^, a small peptide ∼2.7 kDa, in facilitating ASO transport into the CNS.^24^

Tissue and cellular distribution analyses confirmed that ApoB^11^:ASO-α-Syn is actively taken up by neurons in key brain regions, including the substantia nigra, hippocampus, striatum, and cortex, rather than passively diffusing. Additionally, ApoB^11^:ASO -α-Syn uptake by endothelial cells supports receptor-mediated transcytosis across the BBB, while low microglial uptake indicates minimal immune activation, reinforcing its safety profile for long-term use. The absence of significant neuroinflammation is encouraging, especially given the risks associated with chronic neuroinflammatory responses in neurodegenerative disease progression.^25^

Pharmacokinetic and biodistribution data demonstrated rapid systemic availability (early plasma peak), sustained CNS exposure, and pronounced accumulation in the liver and spleen, suggesting dual clearance and therapeutic engagement pathways. Moderate retention in muscle supports the CNS targeted nature of this delivery platform. These findings collectively validate ApoB^11^ as a dual function carrier with both BBB permeability and hepatic uptake, complementing systemic ASO strategies. Our findings also align with previous studies using ApoB-based CNS-targeted therapeutics. For example, ApoB^11^-neprilysin fusion proteins enhanced amyloid-beta clearance via BBB penetration in APPtg mice,^26^ reinforcing the versatility of ApoB-mediated delivery systems for neurodegenerative targets.

One critical observation was the toxicity associated with high-dose (32 mg/kg) ApoB^11^:ASO-α-Syn administration, primarily due to peptide ASO precipitation. The ApoB^11^ peptide is highly hydrophobic, with solubility maintained by its hydrophilic arginine tail; however, ASO binding masks this tail, leading to precipitation.^27^ Upon injection, this aggregation likely caused vascular occlusion, contributing to increased mortality. Thus, peptide:ASO formulation at higher doses presents a challenge for clinical translation. Future formulation improvements and linker chemistries are needed to mitigate aggregation while preserving CNS targeting efficacy.

Additionally, we observed that high-dose treatment caused significant α-Syn knockdown, which may have unintended physiological consequences. α-Syn is known to play critical roles in synaptic vesicle trafficking, neurotransmitter release, and synaptic plasticity._28-30_ Therefore, excessive suppression may disrupt basal neurotransmission or cognitive performance. This concern is supported by studies in α-Syn knockout (Snca⁻/⁻) mice, which show altered dopaminergic signaling, synaptic dysfunction, and deficits in learning and memory.^31-33^ These findings underscore the importance of carefully optimizing the therapeutic window to balance efficacy with potential off-target or compensatory effects. Optimizing the therapeutic window is essential to avoid deleterious effects while maintaining efficacy.

Our time course analysis following a single 2 mg/kg intraperitoneal dose revealed that α-Syn levels dropped significantly by Day 7 (∼30%), peaked at Day 14 (∼60%), and remained suppressed through Day 21, gradually returning to baseline by Day 35. These temporal dynamics, confirmed by qRT-PCR and immunohistochemistry, are consistent with transcriptional knockdown and ASO pharmacokinetics. Importantly, this therapeutic timeline aligns with the observed brain half-life (t₁/₂ = 646.2 hours, or ∼27 days), indicating that target knockdown closely tracks ASO presence in the CNS. This emphasizes the need for careful dosing regimen design to achieve sustained therapeutic benefit without over-suppression or toxicity. Given the prolonged CNS retention in mice, human half-life is likely even longer due to slower clearance in larger species. Allometric scaling, commonly used in ASO development, suggests the potential for monthly or bi-monthly dosing in humans, though this requires validation in primates and clinical studies.^34^ These PK-PD dynamics are essential for optimizing future dosing regimens.

Compared with existing methods like intracerebroventricular (ICV) injection,^2^ our systemic delivery strategy offers a less invasive, more clinically feasible route while still achieving durable CNS retention and α-Syn suppression. This is particularly relevant for clinical translation in chronic diseases. Ligand-mediated transport platforms such as TfR-targeting oligonucleotide transport vehicles (OTVs) have shown similar success,^35^ but exhibit more extensive peripheral distribution due to ubiquitous TfR expression.^36,37^ Similar to TfR, the LDL-receptor is widely expressed on nearly all cells with over 70% of expression in the liver. Other LDL receptor family members are differentially expressed with VLDL and ApoE-receptor expressed in greater abundance in the CNS.^38^ This pattern of expression closely mimics the distribution of the ApoB^11^-ASO-α-Syn we observed. Compared to prior ASO studies in Alzheimer’s disease (AD) and amyotrophic lateral sclerosis (ALS) models,^39,40^ our strategy offers enhanced BBB penetration, non-invasive systemic delivery, and selective neuronal uptake, setting it apart from intrathecal or ICV delivery methods that may hinder patient compliance and clinical scalability.^41^

Several limitations must be addressed. First, while dose-dependent effects are evident, we observed a plateau at 16 mg/kg, suggesting a ceiling effect in α-Syn suppression. Future studies should explore intermediate dosing, improved solubility formulations, and more precise titration to balance efficacy and safety. Second, while our study focused on biochemical outcomes, we did not assess behavioral improvements or long-term neuropathology. These are essential for validating functional benefit.^42^

In conclusion, this study highlights the potential of ApoB^11^:ASO-α-Syn as a promising therapeutic for synucleinopathies. Its ability to cross the BBB and achieve robust knockdown in neurons makes it a strong candidate for further development. With continued optimization of dosage, formulation, and delivery strategies, ApoB^11^ mediated ASO therapies may offer a clinically viable solution for diseases driven by α-Syn pathology.

## MATERIALS and METHODS

### Ethical statements

All animal experiments conducted in this study were reviewed and approved by the UCSD Institutional Animal Care and Use Committee in accordance with the NIH Guide for the Care and Use of Laboratory Animals (protocol #S02221). Measures were taken to minimize discomfort and distress in animals, including the administration of appropriate anesthesia and analgesics during invasive procedures, as well as the implementation of humane endpoints when necessary. Furthermore, the study design adhered to the principles of the 3Rs (Replacement, Reduction, and Refinement) to optimize animal welfare and minimize the number of animals used.

### Animals

Male C57BL/6J mice (RRID:IMSR_JAX:000664) aged 10–12 weeks and weighing 20–30 g were obtained from The Jackson Laboratory for pharmacokinetics and biodistribution studies, while both male and female C57BL/6J mice were used for toxicology assessments to evaluate potential sex-dependent effects. For drug efficacy studies, D-line (PDGFβ-hu-α-Syn) transgenic mice (RRID:IMSR_JAX:038775) were used, which overexpress human α-synuclein (SNCA) under the platelet-derived growth factor B (PDGFB) promoter, leading to progressive α-Syn pathology, synaptic dysfunction, and motor impairments, making them a relevant model for synucleinopathies such as Parkinson’s disease (PD) and dementia with Lewy bodies (DLB).^20,43^ Mice were housed in a specific pathogen-free (SPF) animal facility under controlled conditions (temperature: 24°C ±2°C, relative humidity: 40%–70%, 12-hour light/dark cycle) with *ad libitum* access to standard chow and water. For the toxicology experiment, daily monitoring of body weight and general well-being was conducted to assess potential signs of distress or toxicity. At the study endpoint, mice were euthanized humanely under isoflurane anesthesia, followed by cervical dislocation or cardiac perfusion, in accordance with AVMA (American Veterinary Medical Association) guidelines.

### Preparation of Antisense oligonucleotides, peptide vectors and drug formulation

For this study, we utilized ApoB^11^, a novel peptide derived from the ApoB^38^ peptide, which has been previously shown to facilitate the transport of biomolecules across the BBB.^23,44,45^ ApoB^11^ retains the LDL receptor binding domain of ApoB^38^ (NH2-RLTRKRGLKLAGGGGGRRRRRRRRR) and was synthesized at ≥90% purity.^13,23,46^ For oligonucleotide delivery, a 5-glycine linker and 9-arginine residues were added to its C-terminus, enhancing flexibility and charge-based binding to negatively charged nucleotides. The 2′-OMe-modified ASO-α-Syn (5′-GAC UUU CAA AGG CCA AGG A), targeting nucleotides 168–186 of α-Syn, was selected for its dual specificity to human and mouse α-Syn.^13^ A scrambled ASO control (5′-GGG CAU ACU GAG CUA ACA A) was used for comparison. Both antisense oligonucleotides (ASOs) were synthesized, HPLC-purified, and desalted by Integrated DNA Technology (IDT). The ASOs were reconstituted in RNase-free water at 100 mM and stored in aliquots. To assess biodistribution in different brain regions, a biotinylated version of ApoB^11^-ASO-α-syn was generated (IDT), allowing for visualization, and tracking of drug localization. For *in vivo* experiments, ApoB^11^ and ASO-α-Syn were complexed at a 1:10 molar ratio, based on prior optimization.^13,14^ The complexes were dissolved in a 100 mM Tris-HCl (pH 7.4) and saline solution to ensure stability and proper biological activity before administration.

### ASO Bioassay

A modified version of the ASO detection protocol by Yu et al. was adapted using MesoScale Discovery (MSD) technology for enhanced sensitivity and detection range.^47^ A capture oligonucleotide (5′-CAG CTT GCA TCC TTG GCC TTT GAA AGT C-Bio) with a 3′ biotin tag was synthesized to specifically bind α-Syn ASO, along with a probe oligonucleotide (5′-TGC AAG CTG) modified with a 5′ phosphate group and 3′ biotin tag (Integrated DNA Technologies). Tissue homogenates were diluted to 5 mg/mL (liver, spleen, muscle) or 25 mg/mL (brain) in Hybridization Buffer (60 mM Na₂HPO₄, 0.9 M NaCl, 0.24% Tween-20). Plasma samples were diluted 1:1 with the same buffer before incubation with 0.05 µM capture oligonucleotide at 37°C for 30 minutes with shaking.

MSD streptavidin-coated 96-well plates were blocked with 1% BSA in Hybridization Buffer containing 500 µg/mL sheared salmon sperm DNA, followed by sequential washing and incubation steps. After hybridization, a T4 ligase reaction was performed using the probe oligonucleotide, and plates were incubated for 2 hours. Detection was carried out by adding Streptavidin-SulfoTag (MSD, 1:500 in PBS-T), followed by washing and reading on an MSD reader. ASO concentrations were quantified using a standard curve of α-Syn ASO generated by spiking control tissue homogenates, and data analysis was performed using Excel PK/PD plug-in.^45^

ASO Biodistribution

To assess the presence of intraperitoneally injected antisense oligonucleotide (ASO) in brain tissue, wild-type (WT) C57BL/6 mice (n = 3, male, 10-12 weeks old) were administered biotin-labeled α-Syn ASO bound to the ApoB^11^ peptide at a dose of 2 mg/kg. Mice were perfused transcardially with saline 24 hours post-injection, and brains, liver, muscle, and spleen were collected for analysis. Sagittal brain sections were processed and stained using the Tyramide Red Signal Amplification Kit (AKOYA Biosciences, Cat# SKU NEL702001KT) to enhance the detection of the biotin-tagged ASO.^48^ Central nervous system (CNS) cells were labeled with the following antibodies: anti-NeuN (Millipore, Cat# MAB377, RRID: AB_2298772) for neurons, anti-GFAP (Millipore, Cat# MAB3402, RRID:AB_94844) for astrocytes, anti-Olig2 (Novus, Cat# NBP1-28667, RRID:AB_1914109) for oligodendrocytes, anti-Iba1 (FUJIFILM Wako Pure Chemical Corporation Cat# 019-19741, RRID:AB_839504) for microglia, and Tomato-lectin (Bioworld, Cat# 21761030-1, RRID:AB_2336416) for endothelial cells. These primary antibodies were followed by secondary staining with Alexa Fluor-488 Goat anti-Mouse (Molecular Probes Cat# A-11008, RRID:AB_143165) for NeuN and GFAP, and Alexa Fluor-488 Goat anti-Rabbit (Thermo Fisher Scientific Cat# A-11001, RRID:AB_2534069) for Olig2 and Iba1, for visualization. For the liver, spleen, and muscle tissues, the sections were stained with Wheat germ agglutinin-Alexa Flour-488 and counterstained with DAPI (Hoechst, Invitrogen, Cat# 447167) to highlight nuclear architecture. The sections were then mounted using Prolong Gold anti-fade mounting medium with DAPI (Invitrogen, Cat# P36931) to preserve fluorescence. Imaging was performed using a Leica SP8 LIGHTNING confocal microscope (RRID:SCR_018169), and quantitative image analysis was conducted using ImageJ software (RRID:SCR_003070) to assess ASO distribution and cellular colocalization.

### Pharmacokinetic analysis

The pharmacokinetics of ApoB^11^-α-Syn ASO were assessed in 10-12 week C57BL/6 male mice (n=4 per time point). Mice received a single intraperitoneal (IP) bolus injection of ApoB^11^-α-Syn ASO at 2 mg/kg, and blood plasma was collected at 30 minutes, 1 hour, 2 hours, 4 hours, 8 hours, 16 hours, 24 hours, 48 hours, and 96 hours post-injection. Following plasma collection, mice were saline-perfused and sacrificed, after which tissues including the liver, spleen, quadriceps muscle, and brain were dissected, flash-frozen in liquid nitrogen, and stored at -80°C until further analysis.

Tissues were homogenized using a TissueLyzer in homogenization buffer, 1X nuclease-free PBS containing RNAse inhibitor (NEB) and protease inhibitor (Mini-Complete, MilliporeSigma) using the Bead Mill 24 homogenizer (Thermo Fisher Scientific) with 1.4 mm ceramic beads (Thermo Fisher Scientific). Protein concentrations were determined by BCA assay (Thermo Fisher Scientific), and samples were diluted to 5 mg/mL in ASO hybridization buffer. Each sample was assayed in duplicate using the ASO bioassay, following a 1:2 dilution in hybridization buffer. Assays were repeated at least twice for each tissue type to ensure accuracy. Data analysis and visualization were performed using the Microsoft Excel plug-in pkpdtools.xll, as described by Spencer et al.^26^

### Histopathological Analysis

Mice (wild-type C57BL/6, 10-12 weeks, male/female) were randomly assigned to three groups: control (saline), low-dose (10 mg/kg ApoB^11^:ASO-α-Syn), and high-dose (32 mg/kg ApoB^11^:ASO-α-Syn). Treatments were administered via intraperitoneal injection once a week for four weeks. Animals were euthanized after four weeks of treatment, and major organs, including the liver, kidney, heart, brain, spleen, lungs, gastrointestinal tract, pancreas, thymus, and skeletal muscle, were collected for histopathological evaluation. Tissues were immediately fixed in 10% neutral-buffered formalin for at least 48 hours and subsequently submitted to Comparative Biosciences, Inc. a contract research organization specializing in toxicologic pathology and histopathology for further processing. Histological preparation included paraffin embedding, sectioning (4-5 µm), and hematoxylin and eosin (H&E) staining. A board-certified veterinary pathologist performed a blinded microscopic evaluation of 21 tissue types (Table S1), assessing for inflammation, necrosis, degeneration, fibrosis, vascular changes, and other cellular abnormalities indicative of toxicity. Findings were compared across treatment groups to identify potential dose-dependent effects.

### Protein extraction and Immunoblot analysis

Following the completion of treatments, mice were euthanized, and brains were then carefully extracted, with the right hemisphere immersion-fixed in 4% phosphate-buffered formaldehyde at 4°C for 48 hours, after which they were sectioned sagittally at 40 µm for neuropathological analysis. The left hemisphere was snap-frozen in liquid nitrogen and stored at -80°C for subsequent biochemical investigations.

Subdissection of brains was performed with the aid of a brain-cutting metal matrix (RBMS 200S, Kent Scientific Corporation) to target regions encompassing both the cortex and hippocampus. The region obtained was from approximately 1.0 mm to 3.0 mm lateral to Bregma to ensure inclusion of the desired structures. Tissue samples were homogenized in 1X nuclease-free PBS supplemented with RNase inhibitor (NEB) and protease inhibitor (Mini-Complete, MilliporeSigma) using the Bead Mill 24 homogenizer (Thermo Fisher Scientific) with 1.4 mm ceramic beads. Protein fractions were extracted by incubating homogenates with 10X RIPA buffer (Thermo Fisher Scientific), followed by quantification using a BCA Protein Assay Kit (Thermo Fisher Scientific).

For immunoblotting, 25 µg of total protein per sample was resolved on a 12% Bis-Tris SDS-PAGE gel (Criterion TGX, BIO-RAD) and transferred onto PVDF membranes using the Trans-Blot Turbo Transfer System (BIO-RAD). Membranes were probed sequentially with primary antibodies against total α-synuclein (BD Transduction Laboratories™, Cat# 610787, RRID:AB_398107), phosphorylated α-synuclein at Ser129 [EP1536Y] (Abcam, Cat# ab51253, RRID: AB_869973), and GAPDH (Cell Signaling Technology, Cat# 2118, RRID:AB_561053), followed by HRP-conjugated secondary antibodies (BioRad). Signal detection was performed using an enhanced chemiluminescence (ECL) kit (Thermo Fisher Scientific), and band intensities were quantified with the ChemiDoc Imaging System (Bio-Rad). To ensure consistency in protein normalization, membranes were stripped (Restore Western Blot Stripping Buffer, Thermo Fisher Scientific) and re-probed sequentially in the order of p-Syn, total Syn, and GAPDH. Band intensities were normalized to GAPDH and compared to controls.

### RNA extraction and qPCR

RNA was extracted from the remaining homogenized samples using the RNeasy Kit (Qiagen) following the manufacturer’s instructions. The purity and concentration of RNA were assessed using the Invitrogen Nanodrop One Spectrophotometer. For cDNA synthesis, 500 ng of total RNA was reverse transcribed using the iScript gDNA Clear cDNA Synthesis Kit (BioRad). Real-time PCR (RT-PCR) was performed using the IDT PrimeTime Gene Expression Master Mix (IDT) with primer/probe sets specific for human α-synuclein (IDT, Hs.PT.58.912923). Reactions were carried out using the CFX96 Real-Time PCR System (BioRad). The relative cDNA levels were quantified using the comparative threshold cycle (ΔΔCt) method, with mouse β-actin (IDT, Mm.PT.39a.22214843.g) as an internal control. Expression levels were normalized to wild-type mice treated with ApoB^11^:Scr ASO, and relative mRNA levels were calculated using the previously described 2^(-ΔΔCt) method.^49^

### Immunohistochemistry

Analysis was conducted on free-floating, 40 μm-thick vibratome-cut sections, processed in a blind-coded manner. Sagittal brain sections, taken between 1.5 mm and 2 mm from the midline, were collected, yielding approximately 10-12 slices per mouse. From these, three sections were randomly selected for immunohistochemistry.

For neuropathological assessment, sections were incubated overnight at 4°C with primary antibodies against total α-synuclein (BD Transduction Laboratories, Cat# 610787, RRID: AB_398108) and phosphorylated α-synuclein (Ser129) (Abclonal, Cat# AP0450, RRID: AB_2771549). Following incubation, sections were treated with biotinylated secondary antibodies (Horse anti-mouse IgG (H+L), BA-2000 / Goat anti-rabbit IgG (H+L), BA-1000, Vector Laboratories) and visualized using avidin-biotin complex staining (VECTASTAIN Elite ABC-HRP Kit, Peroxidase-PK-6100) with diaminobenzidine (DAB). High-resolution images were captured using the NanoZoomer S60 Digital Slide Scanner (Hamamatsu, 20X magnification), and regions of interest were identified with NDP.View2 software. Quantification was performed using ImageJ2 (version 2.16.0), with adjusted optical density measurements obtained by averaging values from three distinct areas per region, using corpus callosum background values for normalization.

### Statistical Analysis

All statistical analyses were performed using GraphPad Prism (version 10.0), except for the pharmacokinetic (PK) analysis, which was conducted using the Excel plug-in pkdptools. Data are presented as mean ± standard deviation (SD) with statistical significance set at p < 0.05. For group comparisons, one-way ANOVA followed by Dunnett’s post-hoc test was used when comparing multiple treatment groups against a common control (D-line mice injected with saline). When comparing between individual treatment groups (ASO-α-Syn vs. ASO-Scr), either Tukey-Kramer’s post-hoc test or an unpaired Student’s t-test was applied, depending on the experimental design. For growth curve analysis, two-way ANOVA followed by Tukey’s multiple comparison test was used to assess differences over time.

## DATA AND CODE AVAILABILITY

All data are included in the manuscript. Raw data are available on request.

## ACKNOWLEDGMENTS

The authors thank James Barlow, Floyd Sarsoza, Jazmin Florio, Michael Mante, Estefani Guzman, and Carlos Arias from the Rissman’s lab for technical assistance with this project. This work was supported by NIH/NIA R01 grants AG018440 and AG073979 to RAR and the UCSD Microscopy Core funded by NINDS NS047101. Figures 3A, 4A, and graphical abstract were created with Bio-Render (biorender.com).

## AUTHOR CONTRIBUTIONS

R.U.A., B.S., and R.A.R. conceived the study. R.U.A. and B.S. performed the formal analysis and investigation. B.Q. and S.S. contributed to the investigation. R.U.A. wrote the manuscript.

B.S. and R.A.R. reviewed and edited the manuscript. R.A.R. acquired funding and supervised the work. All authors approved the final version of the manuscript.

## References

1. Zhao, H.T., John, N., Delic, V., Ikeda-Lee, K., Kim, A., Weihofen, A., Swayze, E.E., Kordasiewicz, H.B., West, A.B., and Volpicelli-Daley, L.A. (2017). LRRK2 Antisense Oligonucleotides Ameliorate α-Synuclein Inclusion Formation in a Parkinson’s Disease Mouse Model. Mol Ther Nucleic Acids 8, 508–519. 10.1016/j.omtn.2017.08.002.

2. Cole, T.A., Zhao, H., Collier, T.J., Sandoval, I., Sortwell, C.E., Steece-Collier, K., Daley, B.F., Booms, A., Lipton, J., Welch, M., et al. (2021). α-Synuclein antisense oligonucleotides as a disease-modifying therapy for Parkinson’s disease. JCI Insight 6. 10.1172/jci.insight.135633.

3. Bennett, C.F., and Swayze, E.E. (2010). RNA targeting therapeutics: molecular mechanisms of antisense oligonucleotides as a therapeutic platform. Annu Rev Pharmacol Toxicol 50, 259–293. 10.1146/annurev.pharmtox.010909.105654.

4. Rigo, F., Chun, S.J., Norris, D.A., Hung, G., Lee, S., Matson, J., Fey, R.A., Gaus, H., Hua, Y., Grundy, J.S., et al. (2014). Pharmacology of a central nervous system delivered 2’-O-methoxyethyl-modified survival of motor neuron splicing oligonucleotide in mice and nonhuman primates. J Pharmacol Exp Ther 350, 46–55. 10.1124/jpet.113.212407.

5. Khvorova, A., and Watts, J.K. (2017). The chemical evolution of oligonucleotide therapies of clinical utility. Nat Biotechnol 35, 238–248. 10.1038/nbt.3765.

6. Fletcher, S., Honeyman, K., Fall, A.M., Harding, P.L., Johnsen, R.D., and Wilton, S.D. (2006). Dystrophin expression in the mdx mouse after localised and systemic administration of a morpholino antisense oligonucleotide. J Gene Med 8, 207–216. 10.1002/jgm.838.

7. Hinderer, C., Katz, N., Dyer, C., Goode, T., Johansson, J., Bell, P., Richman, L., Buza, E., and Wilson, J.M. (2020). Translational Feasibility of Lumbar Puncture for Intrathecal AAV Administration. Mol Ther Methods Clin Dev 17, 969–974. 10.1016/j.omtm.2020.04.012.

8. Finkel, R.S., Mercuri, E., Darras, B.T., Connolly, A.M., Kuntz, N.L., Kirschner, J., Chiriboga, C.A., Saito, K., Servais, L., Tizzano, E., et al. (2017). Nusinersen versus Sham Control in Infantile-Onset Spinal Muscular Atrophy. N Engl J Med 377, 1723–1732. 10.1056/NEJMoa1702752.

9. Hua, Y., Sahashi, K., Rigo, F., Hung, G., Horev, G., Bennett, C.F., and Krainer, A.R. (2011). Peripheral SMN restoration is essential for long-term rescue of a severe spinal muscular atrophy mouse model. Nature 478, 123–126. 10.1038/nature10485.

10. Juliano, R.L. (2016). The delivery of therapeutic oligonucleotides. Nucleic Acids Res 44, 6518–6548. 10.1093/nar/gkw236.

11. Min, H.S., Kim, H.J., Naito, M., Ogura, S., Toh, K., Hayashi, K., Kim, B.S., Fukushima, S., Anraku, Y., Miyata, K., and Kataoka, K. (2020). Systemic Brain Delivery of Antisense Oligonucleotides across the Blood-Brain Barrier with a Glucose-Coated Polymeric Nanocarrier. Angew Chem Int Ed Engl 59, 8173–8180. 10.1002/anie.201914751.

12. Soutschek, J., Akinc, A., Bramlage, B., Charisse, K., Constien, R., Donoghue, M., Elbashir, S., Geick, A., Hadwiger, P., Harborth, J., et al. (2004). Therapeutic silencing of an endogenous gene by systemic administration of modified siRNAs. Nature 432, 173–178. 10.1038/nature03121.

13. Spencer, B., Trinh, I., Rockenstein, E., Mante, M., Florio, J., Adame, A., El-Agnaf, O.M.A., Kim, C., Masliah, E., and Rissman, R.A. (2019). Systemic peptide mediated delivery of an siRNA targeting α-syn in the CNS ameliorates the neurodegenerative process in a transgenic model of Lewy body disease. Neurobiol Dis 127, 163–177. 10.1016/j.nbd.2019.03.001.

14. Leitão, A.D.G., Ahammad, R.U., Spencer, B., Wu, C., Masliah, E., and Rissman, R.A. (2023). Novel systemic delivery of a peptide-conjugated antisense oligonucleotide to reduce α-synuclein in a mouse model of Alzheimer’s disease. Neurobiol Dis 186, 106285. 10.1016/j.nbd.2023.106285.

15. Stefanis, L. (2012). α-Synuclein in Parkinson’s disease. Cold Spring Harb Perspect Med 2, a009399. 10.1101/cshperspect.a009399.

16. Morris, H.R., Spillantini, M.G., Sue, C.M., and Williams-Gray, C.H. (2024). The pathogenesis of Parkinson’s disease. Lancet 403, 293–304. 10.1016/s0140-6736(23)01478-2.

17. Games, D., Valera, E., Spencer, B., Rockenstein, E., Mante, M., Adame, A., Patrick, C., Ubhi, K., Nuber, S., Sacayon, P., et al. (2014). Reducing C-terminal-truncated alpha-synuclein by immunotherapy attenuates neurodegeneration and propagation in Parkinson’s disease-like models. J Neurosci 34, 9441–9454. 10.1523/jneurosci.5314-13.2014.

18. Nakamori, M., Junn, E., Mochizuki, H., and Mouradian, M.M. (2019). Nucleic Acid-Based Therapeutics for Parkinson’s Disease. Neurotherapeutics 16, 287–298. 10.1007/s13311-019-00714-7.

19. Crooke, S.T., Baker, B.F., Crooke, R.M., and Liang, X.H. (2021). Antisense technology: an overview and prospectus. Nat Rev Drug Discov 20, 427–453. 10.1038/s41573-021-00162-z.

20. Masliah, E., Rockenstein, E., Veinbergs, I., Mallory, M., Hashimoto, M., Takeda, A., Sagara, Y., Sisk, A., and Mucke, L. (2000). Dopaminergic loss and inclusion body formation in alpha-synuclein mice: implications for neurodegenerative disorders. Science 287, 1265–1269. 10.1126/science.287.5456.1265.

21. Rockenstein, E., Mallory, M., Hashimoto, M., Song, D., Shults, C.W., Lang, I., and Masliah, E. (2002). Differential neuropathological alterations in transgenic mice expressing alpha- synuclein from the platelet-derived growth factor and Thy-1 promoters. J Neurosci Res 68, 568–578. 10.1002/jnr.10231.

22. Kreuter, J., Shamenkov, D., Petrov, V., Ramge, P., Cychutek, K., Koch-Brandt, C., and Alyautdin, R. (2002). Apolipoprotein-mediated transport of nanoparticle-bound drugs across the blood-brain barrier. J Drug Target 10, 317–325. 10.1080/10611860290031877.

23. Masliah, E., and Spencer, B. (2015). Applications of ApoB LDLR-Binding Domain Approach for the Development of CNS-Penetrating Peptides for Alzheimer’s Disease. Methods Mol Biol 1324, 331–337. 10.1007/978-1-4939-2806-4_21.

24. Haque, U.S., and Yokota, T. (2023). Enhancing Antisense Oligonucleotide-Based Therapeutic Delivery with DG9, a Versatile Cell-Penetrating Peptide. Cells 12. 10.3390/cells12192395.

25. Giri, P.M., Banerjee, A., Ghosal, A., and Layek, B. (2024). Neuroinflammation in Neurodegenerative Disorders: Current Knowledge and Therapeutic Implications. Int J Mol Sci 25. 10.3390/ijms25073995.

26. Spencer, B., Verma, I., Desplats, P., Morvinski, D., Rockenstein, E., Adame, A., and Masliah, E. (2014). A neuroprotective brain-penetrating endopeptidase fusion protein ameliorates Alzheimer disease pathology and restores neurogenesis. J Biol Chem 289, 17917–17931. 10.1074/jbc.M114.557439.

27. Gordon, S.M., Pourmousa, M., Sampson, M., Sviridov, D., Islam, R., Perrin, B.S., Jr., Kemeh, G., Pastor, R.W., and Remaley, A.T. (2017). Identification of a novel lipid binding motif in apolipoprotein B by the analysis of hydrophobic cluster domains. Biochim Biophys Acta Biomembr 1859, 135–145. 10.1016/j.bbamem.2016.10.019.

28. Vargas, K.J., Makani, S., Davis, T., Westphal, C.H., Castillo, P.E., and Chandra, S.S. (2014). Synucleins regulate the kinetics of synaptic vesicle endocytosis. J Neurosci 34, 9364–9376. 10.1523/jneurosci.4787-13.2014.

29. Sharma, M., and Burré, J. (2023). α-Synuclein in synaptic function and dysfunction. Trends Neurosci 46, 153–166. 10.1016/j.tins.2022.11.007.

30. Nemani, V.M., Lu, W., Berge, V., Nakamura, K., Onoa, B., Lee, M.K., Chaudhry, F.A., Nicoll, R.A., and Edwards, R.H. (2010). Increased expression of alpha-synuclein reduces neurotransmitter release by inhibiting synaptic vesicle reclustering after endocytosis. Neuron 65, 66–79. 10.1016/j.neuron.2009.12.023.

31. Cabin, D.E., Shimazu, K., Murphy, D., Cole, N.B., Gottschalk, W., McIlwain, K.L., Orrison, B., Chen, A., Ellis, C.E., Paylor, R., et al. (2002). Synaptic vesicle depletion correlates with attenuated synaptic responses to prolonged repetitive stimulation in mice lacking alpha-synuclein. J Neurosci 22, 8797–8807. 10.1523/jneurosci.22-20-08797.2002.

32. Abeliovich, A., Schmitz, Y., Fariñas, I., Choi-Lundberg, D., Ho, W.H., Castillo, P.E., Shinsky, N., Verdugo, J.M., Armanini, M., Ryan, A., et al. (2000). Mice lacking alpha-synuclein display functional deficits in the nigrostriatal dopamine system. Neuron 25, 239–252. 10.1016/s0896-6273(00)80886-7.

33. Al-Wandi, A., Ninkina, N., Millership, S., Williamson, S.J., Jones, P.A., and Buchman, V.L. (2010). Absence of alpha-synuclein affects dopamine metabolism and synaptic markers in the striatum of aging mice. Neurobiol Aging 31, 796–804. 10.1016/j.neurobiolaging.2008.11.001.

34. Geary, R.S., Norris, D., Yu, R., and Bennett, C.F. (2015). Pharmacokinetics, biodistribution and cell uptake of antisense oligonucleotides. Adv Drug Deliv Rev 87, 46–51. 10.1016/j.addr.2015.01.008.

35. Barker, S.J., Thayer, M.B., Kim, C., Tatarakis, D., Simon, M.J., Dial, R., Nilewski, L., Wells, R.C., Zhou, Y., Afetian, M., et al. (2024). Targeting the transferrin receptor to transport antisense oligonucleotides across the mammalian blood-brain barrier. Sci Transl Med 16, eadi2245. 10.1126/scitranslmed.adi2245.

36. Carrilho, P., Chopra, S., Pichika, M.R., Fleming, R.E., and Parrow, N.L. (2025). Editorial: The potential of transferrin as a drug target and drug delivery system. Front Pharmacol 16, 1584190. 10.3389/fphar.2025.1584190.

37. Dehouck, B., Fenart, L., Dehouck, M.P., Pierce, A., Torpier, G., and Cecchelli, R. (1997). A new function for the LDL receptor: transcytosis of LDL across the blood-brain barrier. J Cell Biol 138, 877–889. 10.1083/jcb.138.4.877.

38. Beffert, U., Stolt, P.C., and Herz, J. (2004). Functions of lipoprotein receptors in neurons. J Lipid Res 45, 403–409. 10.1194/jlr.R300017-JLR200.

39. Bennett, C.F., Krainer, A.R., and Cleveland, D.W. (2019). Antisense Oligonucleotide Therapies for Neurodegenerative Diseases. Annu Rev Neurosci 42, 385–406. 10.1146/annurev-neuro-070918-050501.

40. Martier, R., Liefhebber, J.M., García-Osta, A., Miniarikova, J., Cuadrado-Tejedor, M., Espelosin, M., Ursua, S., Petry, H., van Deventer, S.J., Evers, M.M., and Konstantinova, P. (2019). Targeting RNA-Mediated Toxicity in C9orf72 ALS and/or FTD by RNAi-Based Gene Therapy. Mol Ther Nucleic Acids 16, 26–37. 10.1016/j.omtn.2019.02.001.

41. Wu, D., Chen, Q., Chen, X., Han, F., Chen, Z., and Wang, Y. (2023). The blood-brain barrier: structure, regulation, and drug delivery. Signal Transduct Target Ther 8, 217. 10.1038/s41392-023-01481-w.

42. Ishimoto, T., Oono, M., Kaji, S., Ayaki, T., Nishida, K., Funakawa, I., Maki, T., Matsuzawa, S.I., Takahashi, R., and Yamakado, H. (2024). A novel mouse model for investigating α- synuclein aggregates in oligodendrocytes: implications for the glial cytoplasmic inclusions in multiple system atrophy. Mol Brain 17, 28. 10.1186/s13041-024-01104-7.

43. Hashimoto, M., Rockenstein, E., and Masliah, E. (2003). Transgenic models of alpha-synuclein pathology: past, present, and future. Ann N Y Acad Sci 991, 171–188.

44. Spencer, B., Marr, R.A., Gindi, R., Potkar, R., Michael, S., Adame, A., Rockenstein, E., Verma, I.M., and Masliah, E. (2011). Peripheral delivery of a CNS targeted, metalo- protease reduces aβ toxicity in a mouse model of Alzheimer’s disease. PLoS One 6, e16575. 10.1371/journal.pone.0016575.

45. Spencer, B., Valera, E., Rockenstein, E., Trejo-Morales, M., Adame, A., and Masliah, E. (2015). A brain-targeted, modified neurosin (kallikrein-6) reduces α-synuclein accumulation in a mouse model of multiple system atrophy. Mol Neurodegener 10, 48. 10.1186/s13024-015-0043-6.

46. Spencer, B., Trinh, I., Rockenstein, E., Mante, M., Florio, J., Adame, A., El-Agnaf, O.M.A., Kim, C., Masliah, E., and Rissman, R.A. (2024). Corrigendum to “Systemic peptide mediated delivery of an siRNA targeting α-syn in the CNS ameliorates the neurodegenerative process in a transgenic model of Lewy Body Disease” [Neurobiology of Disease127 (2019) 163-177]. Neurobiol Dis 191, 106397. 10.1016/j.nbd.2023.106397.

47. Yu, R.Z., Kim, T.W., Hong, A., Watanabe, T.A., Gaus, H.J., and Geary, R.S. (2007). Cross- species pharmacokinetic comparison from mouse to man of a second-generation antisense oligonucleotide, ISIS 301012, targeting human apolipoprotein B-100. Drug Metab Dispos *35*, 460-468. 10.1124/dmd.106.012401.

48. Bobrow, M.N., Harris, T.D., Shaughnessy, K.J., and Litt, G.J. (1989). Catalyzed reporter deposition, a novel method of signal amplification. Application to immunoassays. J Immunol Methods 125, 279–285. 10.1016/0022-1759(89)90104-x.

49. Rao, X., Huang, X., Zhou, Z., and Lin, X. (2013). An improvement of the 2^(-delta delta CT) method for quantitative real-time polymerase chain reaction data analysis. Biostat Bioinforma Biomath 3, 71–85.

