## Supplemental Tables & Figures for "Anti-Sense Oligonucleotide as a Therapeutic for Synucleinopathies: Pharmacokinetic, Safety and Efficacy Evaluation"

***Corresponding Author:**

**Short Title:**

ASO Therapy for Synucleinopathies

**Table S1.** Mortality assessment following high-dose ApoB$^{11}$:ASO-α-syn administration. Survival was monitored following weekly intraperitoneal injections of ApoB$^{11}$:ASO-α-syn at escalating doses (75, 50, and 32 mg/kg). The highest non-lethal dose was determined to be 32 mg/kg, with no mortality observed over the 4-week treatment period.

| ASO-α-syn (mg/kg) | Mice (n) | Mortality (%) | Notes |
|---|---|---|---|
| 75 | 4 | 100% | All dead after first injection |
| 50 | 4 | 25% (1/4) after first injection, 75% (3/4) after 1 week, 100% (4/4) after third dose | Mortality increased over time |
| 32 | 4 | 0% | All survived after 4 doses (weekly) |

**Table S2:** Histopathological Evaluation Tissue list

| Tissue | Control (Saline) | | $ApoB^{11}$:ASO (32 mg/kg) | |
|---|---|---|---|---|
| | M (n=4) | F (n=4) | M (n=4) | F (n=4) |
| Liver | N | N | N | N |
| Esophagus | N | N | N | N |
| Pancrease | N | N | N | N |
| Kidney | N | N | N | N |
| Heart | N | N | N | N |
| Spleen | N | N | N | N |
| Thymus | N | N | N | N |
| Lung | N | N | N | N |
| Tongue | N | N | N | N |
| Small Intestine | N | N | N | N |
| Stomach | N | N | N | N |
| Colon/cecum | N | N | N | N |
| Brain | N | N | N | N |
| Eye/optic nerve | N | N | N | N |
| Harderian Gland | N | N | N | N |
| Skeletal Muscle | N | N | N | N |
| Lymph Node | N | N | N | N |
| Bone | N | N | N | N |
| Bone Marrow | N | N | N | N |
| Cartilage | N | N | N | N |
| Spinal Cord | N | N | N | N |
| Gall Blader | N | N | N | N |
| Salivary Gland | N | N | N | N |

N= No lesion observed

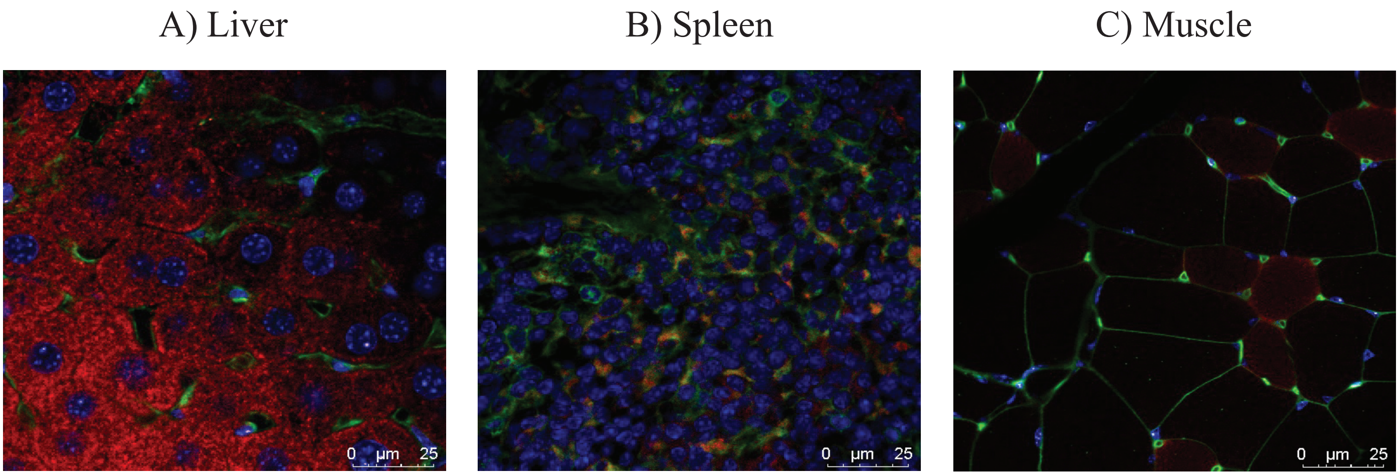


**Figure S1**. Peripheral tissue distribution of ApoB$^{11}$:ASO-α-syn following systemic delivery. (A–C) Representative immunofluorescence images of liver, spleen, and skeletal muscle collected 24 hours post intraperitoneal injection of biotin-labeled ApoB$^{11}$:ASO-α-syn (2 mg/kg) in wild-type C57BL/6 mice (n = 3). Images show ASO localization across different tissue compartments. Scale bar = 50 µm.

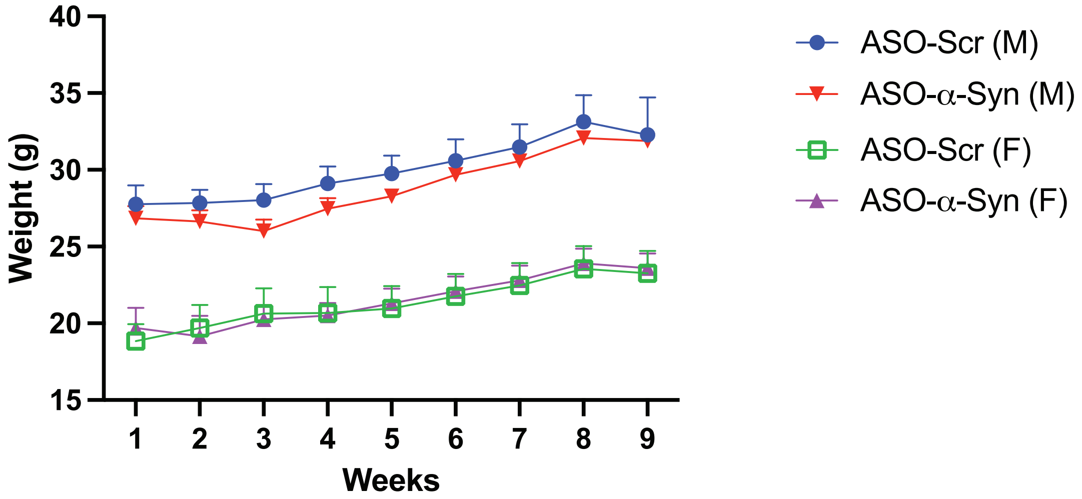


**Figure S2:** Body weight monitoring following high-dose ApoB[11]:ASO-α-syn treatment. Weekly administration of 32 mg/kg ApoB[11]:ASO-α-syn over four weeks, followed by a four-week recovery period, did not cause significant changes in body weight or overall health relative to saline-treated controls.
